# Evaluation of Reproducible and Transparent Research Practices in Pulmonology Publications

**DOI:** 10.1101/726505

**Authors:** Caleb A. Smith, Johnny Nolan, Daniel J. Tritz, Trace E. Heavener, Jameson Pelton, Kathy Cook, Matt Vassar

## Abstract

**Rationale:** Study reproducibility is valuable for validating or refuting results. Provision of reproducibility indicators, such as materials, protocols, and raw data in a study to improve its potential for reproduction. Efforts to reproduce noteworthy studies in the biomedical sciences have resulted in an overwhelming majority of them being found to be unreplicable, causing concern for the integrity of research in other fields, including medical specialities.

**Objective:** Here, we analyzed the reproducibility of studies in the field of pulmonology.

**Methods:** 300 pulmonology articles were randomly selected from an initial PubMed search for data extraction. Two authors scoured these articles for reproducibility indicators including materials, protocols, raw data, analysis scripts, inclusion in systematic reviews, and citations by replication studies as well as other factors of research transparency including open accessibility, funding source and competing interest disclosures, and study preregistration.

**Main Results:** Few publications included statements regarding materials (11%), protocols (1%), data (21%), and analysis script (0%) availability. Less than 10% indicated preregistration. More than half of the publications analyzed failed to provide a funding statement. Conversely, 66% of the publications were open access and 70% included a conflict of interest statement.

**Conclusion:** Overall, our study indicates pulmonology research is currently lacking in efforts to increase replicability. Future studies should focus on providing sufficient information regarding materials, protocols, raw data, and analysis scripts, among other indicators, for the sake of clinical decisions that depend on replicable or refutable results from the primary literature.

## Introduction

Reproducibility—the ability to duplicate a study’s results using the same materials and methods as the original investigator—is central to the scientific method(1). Study reproducibility establishes confidence in the efficacy of therapies, while results that contradict original findings may lead to overturning previous standards. Herrera-Perez et al. recently evaluated 396 medical reversals in which suboptimal clinical practices were overturned when randomized controlled trials yielded results contrary to current practices(2). Given the evolving nature of evidence-based patient care, studies must be conducted in a way that fosters reproducibility and transparency. Further, materials, protocols, analysis scripts, and patient data must be made available to enable verification.

Efforts supporting reproducibility are becoming more widespread owing to the open science movement. In 2013, the Center for Open Science was established to “increase the openness, integrity, and reproducibility of scientific research”(3). The center sponsored two large-scale reproducibility efforts: a series of 100 replication attempts in psychology and a series of 50 landmark cancer biology study replication attempts. In the first, investigators successfully reproduced only 39% of the original study findings(4). In the second, efforts were halted after only 18 replications because of lack of information and materials from authors, insufficient funding, and insufficient time to perform all the experiments(5). The center also created the Open Science Framework, a repository in which authors may deposit study protocols, participant data, analysis scripts, and other materials needed for study reproduction. More recently, the center created Transparency and Openness Promotion Guidelines, which include eight transparency standards and provides guidance for funders and journals, and initiated the use of badges for journals that adopt reproducible practices.

Current estimates of study reproducibility are alarming. In the biomedical sciences, reproducibility rates may be as low as 25%(6). One survey of 1576 scientists found that 90% of respondents believed science was experiencing a reproducibility crisis; 70% reported not being able to reproduce another investigator’s findings, and more than half reported an inability to reproduce their own findings(7). The picture is even less clear in the clinical sciences. Ioannidis found that of 49 highly cited original research publications, seven were refuted by newer studies, and seven suggested higher efficacy than follow-up results; only 22 were successfully replicated(8). The National Institutes of Health and the National Science Foundation have responded to this crisis by taking measures to ensure that studies funded by tax dollars are more reproducible. However, little is known about the extent to which reproducibility practices are used in clinical research.

In this study, we evaluated reproducible and transparent research practices in the pulmonology literature(9). Our goals were (i) to determine areas of strength and weakness in current use of reproducible and transparent research practices and (ii) to establish a baseline for subsequent investigations of the pulmonology literature.

## Methods

This observational study employed a cross-sectional design. We used the methodology of Hardwicke et al.(9), with modifications. In reporting this study, we follow the guidelines for meta-epidemiological methodology research(10) and the Preferred Reporting Items for Systematic Reviews and Meta-analyses (PRISMA)(11). This study did not satisfy the regulatory definition for human subjects research as specified in the Code of Federal Regulations and therefore was not subject to institutional review board oversight. We have listed our protocol, materials, and data on Open Science Framework (https://osf.io/n4yh5/).

### Journal and Publication Selection

The National Library of Medicine catalog was searched by DT using the subject terms tag “Pulmonary Medicine[ST]” to identify pulmonary medicine journals on May 29, 2019. To meet inclusion criteria, journals had to be published in English and be MEDLINE indexed. We obtained the electronic ISSN (or linking ISSN) for each journal in the NLM catalog meeting inclusion criteria. Using these ISSNs, we formulated a search string and searched PubMed on May 31, 2019, to locate publications published between January 1, 2014, to December 31, 2018. We then randomly selected 300 publications for data extraction using Excel’s random number function (https://osf.io/zxjd9/).

### Extraction Training

Prior to data extraction, two investigators (JN, CS) underwent training to ensure inter-rater reliability. The training included an in-person session to review the study design, protocol, Google form, and location of the extracted data elements in two publications. The investigators were next provided with three additional publications from which to extract data. Afterward, the pair reconciled differences by discussion. This training session was recorded and deposited online for reference (https://osf.io/tf7nw/). Prior to extracting data from all 300 publications, the two investigators extracted data from the first 10, followed by a final consensus meeting. Data extraction for the remaining 290 publications followed, and a final consensus meeting was held to resolve disagreements. A third author (DT) was available for adjudication, if necessary.

### Data Extraction

The two investigators extracted data from the 300 publications in a duplicate and blinded fashion. A pilot-tested Google form was created from Hardwicke et al.(9), with additions (see Table 1 for a description of the indicators of reproducibility and transparency). This form prompted coders to identify whether a study had important information that needed to be reproducible (https://osf.io/3nfa5/). The extracted data varied by study design. Studies without empirical data (e.g., editorials, commentaries [without reanalysis], simulations, news, reviews, and poems) had only the publication characteristics, conflict of interest statement, financial disclosure statement, funding sources, and open access availability. We catalogued the most recent year and 5-year impact factor of the publishing journals. We also included the following study designs: cohort, case series, secondary analysis, chart review, and cross-sectional. Finally, we expanded the funding options to include university, hospital, public, private/industry, or nonprofit.

**Table 1:**
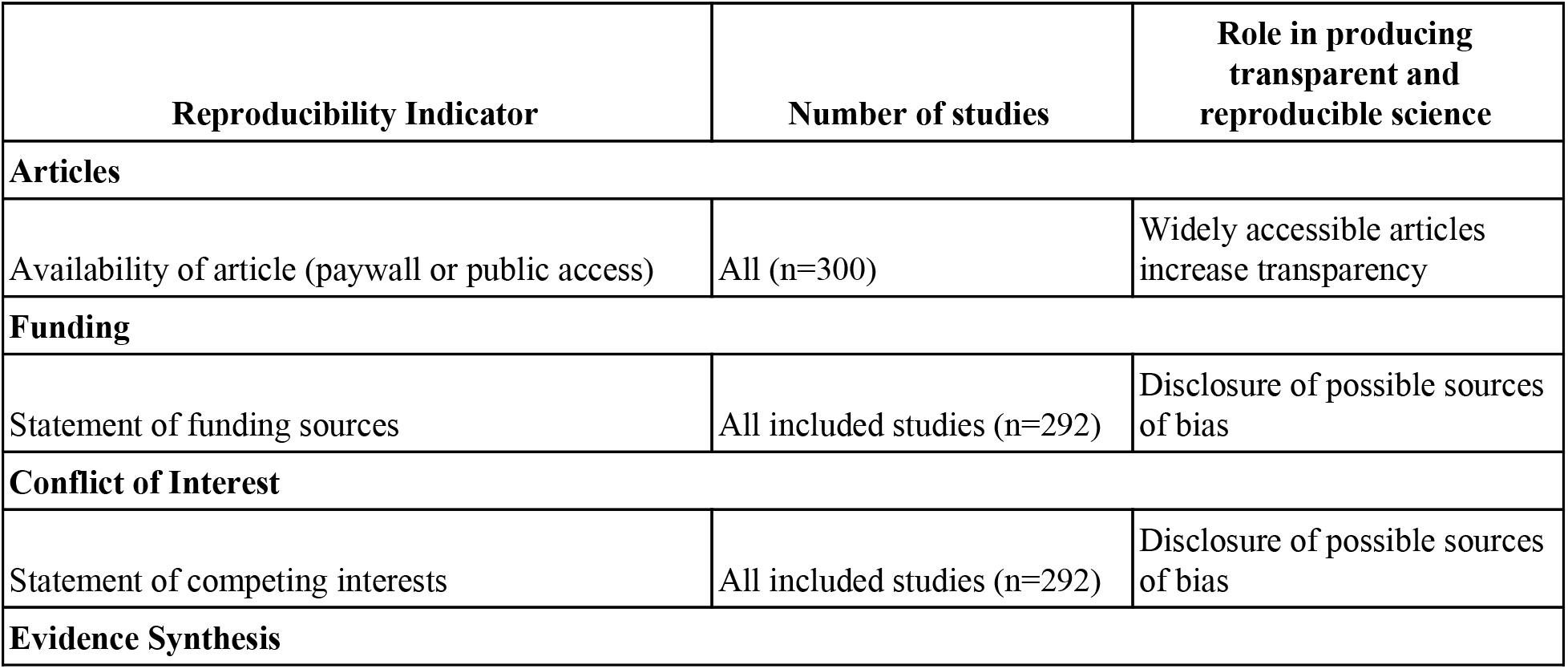
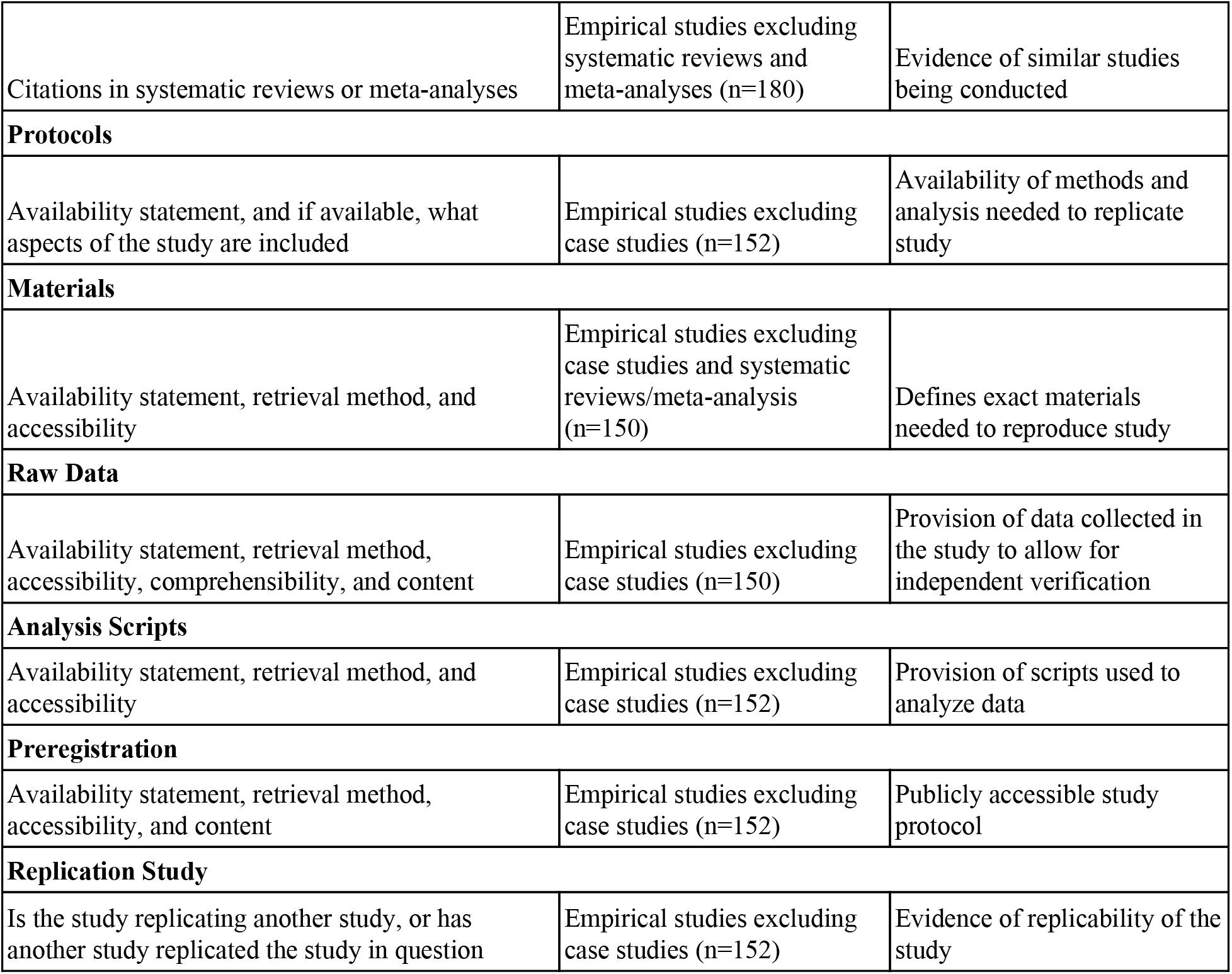
Indicators of Reproducibility

### Verification of Open Access Status of Publications

We used Open Access Button (http://www.openaccessbutton.org) to identify publications as being publicly available. Both the journal title and DOI were used in the search to mitigate chances of missing an article. If Open Access Button could not locate an article, we searched Google and PubMed to confirm open access status.

### Publication Citations Included in Research Synthesis and Replication

For empirical studies, Web of Science was used to identify whether the publication was replicated in other studies and had been included in systematic reviews and/or meta-analyses. To accomplish these tasks, two investigators (CS, JN) inspected the titles, abstracts, and introductions of all publications in which the reference study was cited. This process was conducted in a duplicate, blinded fashion.

### Data Analysis

We used Microsoft Excel to calculate descriptive statistics and 95% confidence intervals (95% CIs).

## Results

### Study Selection and Article Accessibility

Our PubMed search identified 299,255 publications. Limiting our search to articles published from January 1, 2014, to December 31, 2018, yielded 72,579 publications, from which 300 were randomly selected. Of these 300 publications, 194 were open access and 106 were behind a paywall. Eight publications could not be accessed by investigators and were thus excluded, leaving 292 for further analysis (Figure 1). Characteristics of the included publications can be found in Table 2.

**Figure 1:**
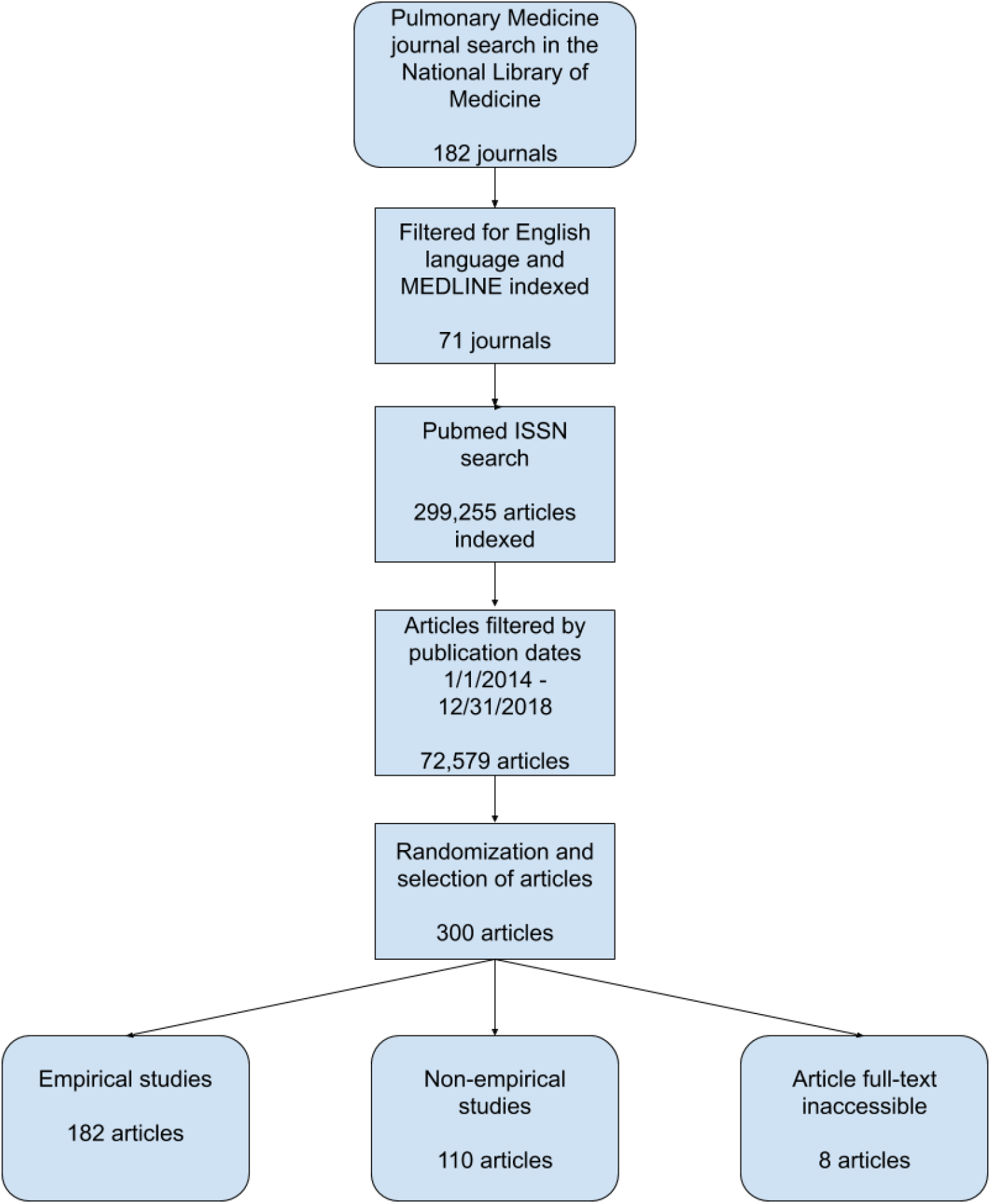
Article selection and filtering process

**Table 2:**
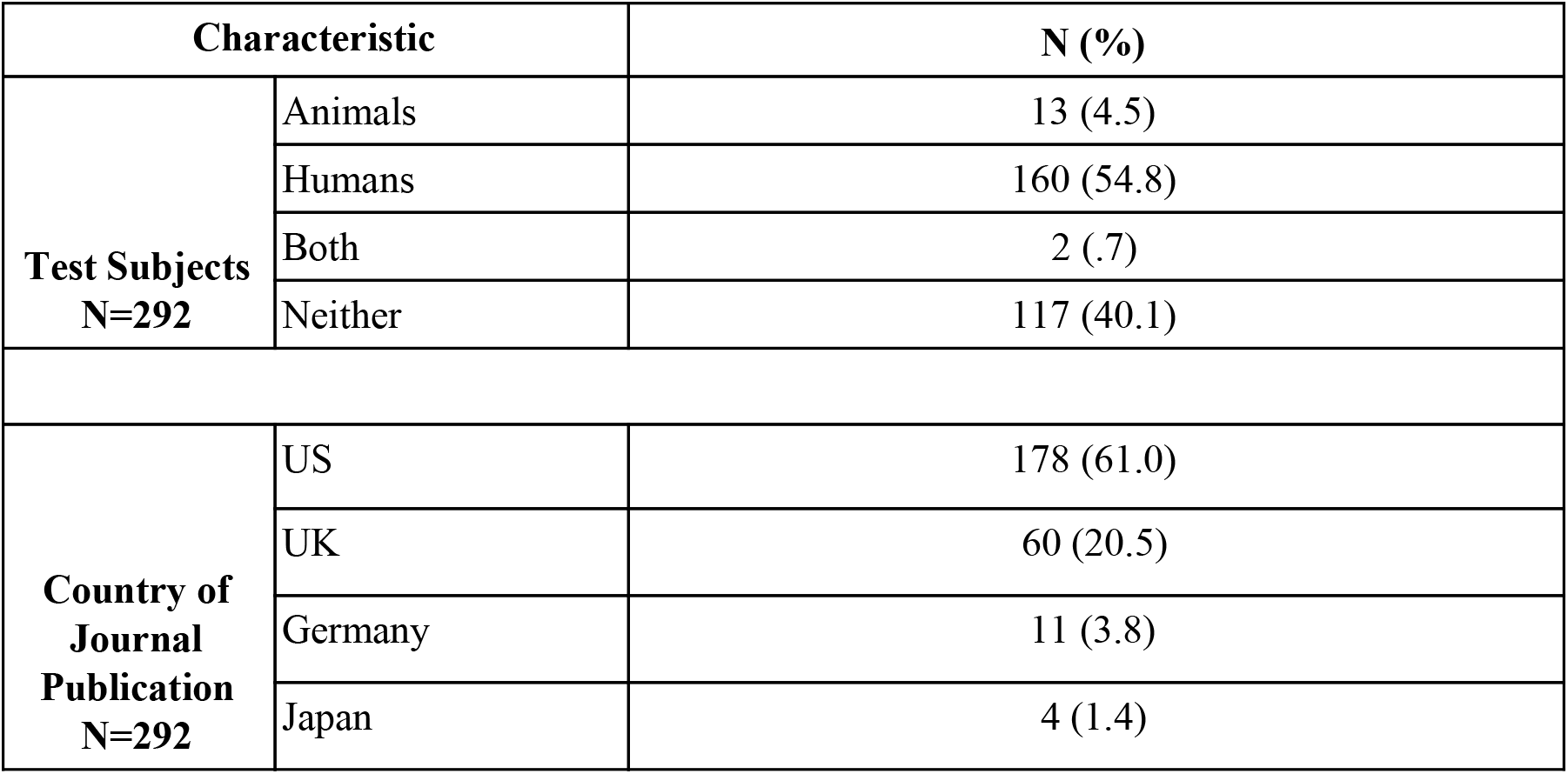
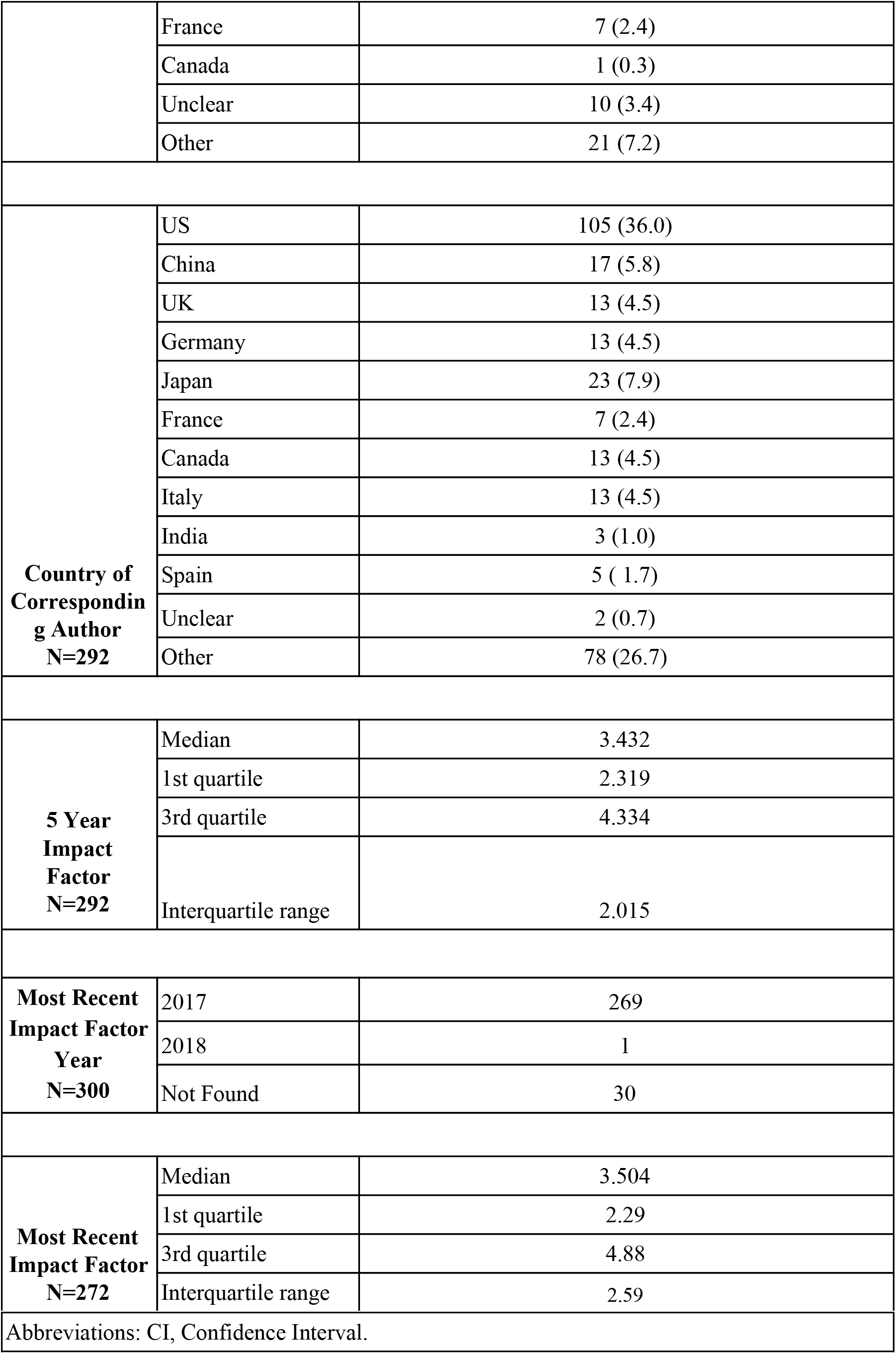
Characteristics of Included Publications

### Availability of Reproducibility Indicators

Figure 2 depicts an overview of our study results. A total of 150 empirical studies (excluding 30 case studies/case series and two meta-analyses) were evaluated for material availability. The majority of studies offered no statement regarding availability of materials (*n* = 133; 88.67% [95% CI, 85.08%–92.25%]). Sixteen studies (10.67% [7.17%–14.16%]) had a clear statement regarding the availability of study materials. One study (0.67% [0%–1.59%]) included an explicit statement that the materials were not publicly available. Ten of the 16 materials files were accessible; the remaining four either led to a broken URL link or a pay-walled request form.

**Figure 2:**
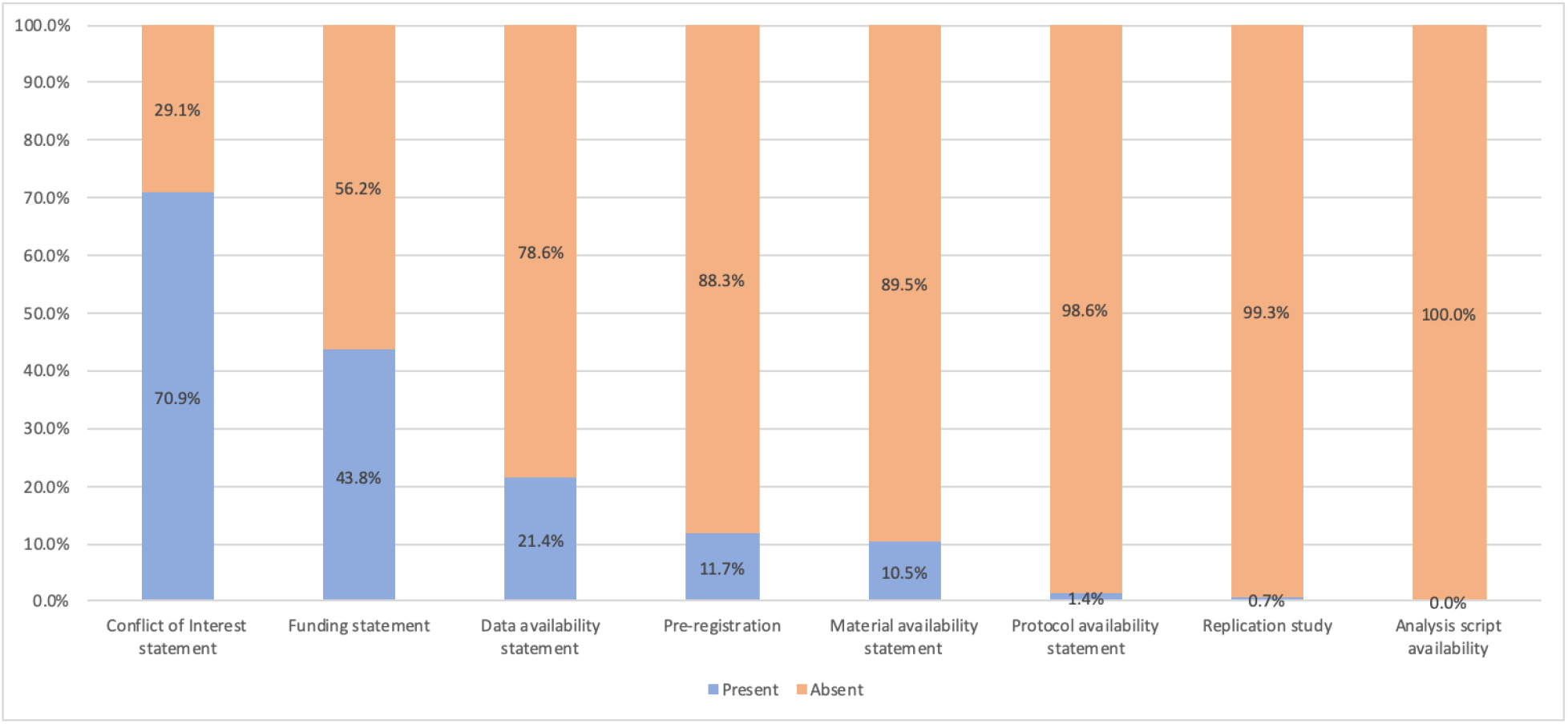
Proportion of Studies with Reproducibility Indicators

A total of 152 empirical studies (excluding 30 case studies/case series) were assessed for availability of protocols, raw data, and analysis scripts. Two studies provided access to a protocol (1.32% [0.03%–2.61%]). Data availability statements were more common, with 31 studies (20.39% [15.84%–24.95%]) including a statement that at least partial data were available. Analysis scripts were not found in a single study. More information on these metrics is presented in Supplemental Table 1.

### Study Preregistration

A total of 152 empirical studies (excluding 30 case studies/case series) were searched for a statement regarding study preregistration. Few studies included statements: 14 (9.21% [5.94%–12.48%]) declared preregistration, while three (1.97% [0.40%–3.55%]) explicitly disclosed that they were not pre-registered. More information on preregistration is presented in Supplemental Table 1.

### Study Replication and Citation Analysis

Of 152 empirical studies analyzed, only one (0.66% [0%–1.57%]) reported replication of the methods of a previously published study. No studies were cited by a replication study. A total of 180 of the 182 empirical studies (excluding two meta-analyses) were evaluated to determine whether any had been included in a systematic review. Thirteen studies (7.22% [4.29%–10.15%]) were cited once in a systematic review or meta-analysis, 11 studies (6.11% [3.40%–8.82%]) were cited in two to five systematic reviews or meta-analyses, and one study (0.56% [0%–1.40%]) was cited in more than five systematic reviews or meta-analyses. One study (0.56% [0.55%–0.55%]) was explicitly excluded from a systematic review.

### Conflict of Interest and Funding Disclosures

All 292 publications were assessed for their inclusion of a conflict of interest statement and/or a funding statement. A majority (*n* = 207; 70.89%) included a conflict of interest statement, with 157 declaring no competing interests (53.77% [48.13%–59.41%]). More than half of the publications failed to provide a funding statement (*n* = 164; 56.16%; Table 3). In publications with a funding statement, public funding was the most common source (*n* = 37; 12.67%).

**Table 3:** Indicators of Reproducibility in Pulmonology Studies

| **Characteristics** |  | **N (%)** | **95% CI** |
| --- | --- | --- | --- |
| **Funding N=292** | University | 6 (2.1) | 0.5-3.7 |
|  | Hospital | 3 (1.0) | 0-2.2 |
|  | Public | 37 (12.7) | 8.9-16.4 |
|  | Private/Industry | 18 (6.2) | 3.4-8.9 |
|  | Non-profit | 5 (1.7) | 0.2-3.2 |
|  | No statement listed | 164 (56.2) | 50.6-61.8 |
|  | Multiple sources | 24 (8.2) | 5.1-11.3 |
|  | No funding received | 35 (12.0) | 8.3-15.7 |
| **Conflict of Interest Statement N=292** | Statement, one or more conflicts of interest | 50 (17.1) | 12.9-21.4 |
|  | Statement, no conflict of interest | 157 (53.8) | 48.1-59.4 |
|  | No conflict of interest statement | 85 (29.1) | 24.0-34.3 |
| **Data Availability N=152** | Statement, some data are available | 50 (17.1) | 12.9-21.4 |
|  | Statement, data are not available | 157 (53.8) | 48.1-59.4 |
|  | No data availability statement | 85 (29.1) | 24.0-34.3 |
| **Material Availability N=150** | Statement, some materials are available | 16 (10.7) | 7.17-14.2 |
|  | Statement, materials are not available | 1 (0.7) | 0-1.6 |
|  | No materials availability statement | 133 (88.7) | 85.1-92.3 |
| **Protocol Availability**<br>**N=152** | Full Protocol | 2 (1.3) | 0-2.6 |
|  | No Protocol | 150 (98.7) | 97.4-100 |
| **Analysis Scripts**<br>**N=152** | Statement, some analysis scripts are available | 0 | 0 |
|  | Statement, analysis scripts are not available | 0 | 0 |
|  | No analysis script availability statement | 152 (100) | 0 |
| **Replication Studies** **N=152** | Novel study | 151 (99.3) | 98.4-100 |
|  | Replication | 1 (0.7) | 0-1.6 |
| **Cited by Systematic Review or Meta-Analysis**<br>**N=180** | No Citations | 155 (86.1) | 82.2-90.0 |
|  | A Single Citation | 13 (7.2) | 4.3-10.2 |
|  | One to Five Citations | 11 (6.1) | 3.4-8.8 |
|  | Greater than Five Citations | 1 (.56) | 0-1.4 |
|  | Excluded in SR or MA | 1 (.56) | 0-1.4 |
| **Preregistration**<br>**N=152** | Statement present, preregistered | 14 (9.7) | 5.9-12.5 |
|  | Statement present, not pre-registered | 3 (2.1) | 0.4-3.5 |
|  | No preregistration statement | 135 (88.8) | 85.2-92.4 |
| **Frequency of Reproducibility Indicators in Selected Studies**<br>**N=292** | **Number of Indicators Present in Study** |  |  |
|  | 0 | 66 (22.6) | — |
|  | 1 | 131(44.9) | — |
|  | 2-5 | 95 (32.5) | — |
|  | 6-8 | 0 | — |
| **Open Access**<br>**N=292** | Found via Open Access Button | 114 | — |
|  | Yes found article via other means | 80 | — |
|  | Could not access through paywall | 98 | — |

## Discussion

In this cross-sectional review of pulmonology publications, a substantial majority failed to provide materials, participant data, or analysis scripts. Many were not preregistered and few had an available protocol. Reproducibility has been viewed as an increasingly troublesome area of study methodology(12). Recent attempts at reproducing preclinical(13, 14) and clinical studies have found that only 25%–61% of studies may be successfully reproduced(6, 15). Many factors contribute to limited study reproducibility, including poor (or limited) reporting of study methodology, prevalence of exaggerated statements, and limited training on experimental design in higher education(16). In an effort to limit printed pages and increase readability, journals may request that authors abridge methods sections(17). Here, we briefly comment on selected indicators to present a balanced view of the perspectives of those in favor of reproducibility and transparency and those who resist enacting such changes.

First, data sharing allows for the independent verification of study results or reuse of that data for subsequent analyses. Two sets of principles exist. The first, known as FAIR, outlines mechanisms for findability, accessibility, interoperability, and reusability. FAIR principles are intended to apply to study data as well as the algorithms, tools, and workflows that led to the data. FAIR advocates that data be accessible to the right people, in the right way, and at the right time(18). A second set of principles relate to making data available to the public for access, use, and share without licenses, copyrights, or patents(19). While we advocate for data sharing, we recognize that it is a complex issue. First, the process for making data available for others’ use requires skills. Further, the process, which includes the construction of data dictionaries and data curation, is time consuming. Furthermore, concerns exist with regard to unrestricted access to data facilitating a culture of “research parasites,” a term coined by Drazen and Longo(20) that suggests that secondary researchers might exploit primary research data for publication. Drazen and Longo also cautioned that secondary authors might not understand the decisions made when defining parameters of the original investigations. Finally, the sensitive nature of some data causes concern among researchers.

Second, preregistering a study requires authors to provide their preliminary protocol, materials, and analysis plan in a publicly available website. The most common websites used by authors are ClinicalTrials.gov and the International Clinical Trial Registry Platform hosted by the World Health Organization. These registries improve the reliability and transparency of published findings by preventing selective reporting of results, preventing unnecessary duplication of studies, and providing relevant material to patients that may enroll in such trials(21). The Food and Drug Administration (FDA) Amendments Act and the International Committee of Medical Journal Editors (ICMJE) have both required registration of clinical trials prior to initiation of a study(22, 23). Selective reporting bias, which includes demoting primary endpoints, omitting endpoints, or upgrading secondary endpoints in favor of statistical significance, may be especially pervasive and problematic. Numerous studies across several fields of medicine have evaluated the extent and magnitude of the problem (24–26). The consequences of selective reporting bias and manipulation of endpoints may compromise clinical decision making. Another issue—*p*-hacking—occurs when researchers repeatedly analyze study data until they achieve statistically significant results. Preregistration of protocols and statistical analysis plans can be used to fact check published papers to ensure that any alterations made in the interim were made for good reason.

Third, transparency related to study funding and financial conflicts of interest should be emphasized. In a previous study, we found that one-third of the authors of pivotal oncology trials underlying FDA drug approvals failed to adequately disclose personal payments from the drug sponsor(27). Recent news accounts of a prominent breast cancer researcher who failed to disclose financial relationships with pharmaceutical companies in dozens of publications has heightened awareness of the pervasiveness of this issue(28). The ICMJE considers willful nondisclosure of financial interests to be a form of research misconduct(29). It is critical that the public be able to adequately evaluate financial relationships of the authors of the published studies in order to evaluate the likelihood of biased results and conclusions.

Several changes are needed to establish a culture of reproducibility and transparency. First, increased awareness of and training about these issues are needed. The National Institutes of Health has funded researchers to produce training and materials, which are available on the Rigor and Reproducibility Initiative website(30), but more remains to be done. Strong mentorship is necessary to encourage trainees to adopt and incorporate reproducible research practices. Research on mentorship programs has found that trainees who have mentors report greater satisfaction with time allocation at work and increased academic self-efficacy compared with trainees without a mentor(31). Conversely, poor mentorship can reinforce poor research practices among junior researchers, such as altering data to produce positive results or changing how results are reported(32). Other research stakeholders must be involved as well. Although many journals recommend the use of reporting guidelines for various study designs, such as CONSORT and PRISMA, evidence suggests that these guidelines are not followed by authors or enforced by journals(33). When journals enforce adherence to reporting guidelines, the completeness of reporting is improved(34). Detractors of reporting guidelines are concerned that certain checklists (CONSTORT, STROBE, STARD) will be used to judge research quality rather than improve writing clarity, that editors and peer reviewers will fail to enforce these guidelines, and that insufficient research exists to evaluate the outcomes from applying these guidelines(35).

Our study has both strengths and limitations. We randomly sampled a large number of pulmonology journals containing various types of publications to generalize our findings across the specialty. Our study design also used rigorous training sessions and a standardized protocol to increase the reliability of our results. In particular, our data extraction process, which involved blinded and duplicate extraction by two investigators, is the gold standard systematic review methodology and is recommended by the Cochrane Collaboration(36). We have made all study materials available for public review to enhance the reproducibility of this study. Regarding limitations, our inclusion criteria for journals (i.e., published in English and MEDLINE indexed) potentially removed journals that contained more lax recommendations regarding indicators of reproducibility and transparency. Furthermore, although we obtained a random sample of publications for analysis, our sample may not have been representative of all pulmonology publications. Our results should be interpreted in light of these strengths and limitations.

In conclusion, our study of the pulmonology literature found that reproducible and transparent research practices are not being incorporated into research. Sharing of study artifacts, in particular, needs improvement. The pulmonology research community should seek to establish norms of reproducible and transparent research practices.

## Author Contributions

DJT, MV: Substantial contributions to the conception and design of the work. CAS, JN, DJT: Acquisition, analysis, and interpretation of data for the work. CAS, DJT, TEH, JP, KC: Drafted the work and revised it critically for important intellectual content. MV: Final approval of the version submitted for publication. CAS: Accountability for all aspects of the work in ensuring that questions related to the accuracy or integrity of any part of the work are appropriately investigated and resolved.

---

**Supplemental Table 1:**
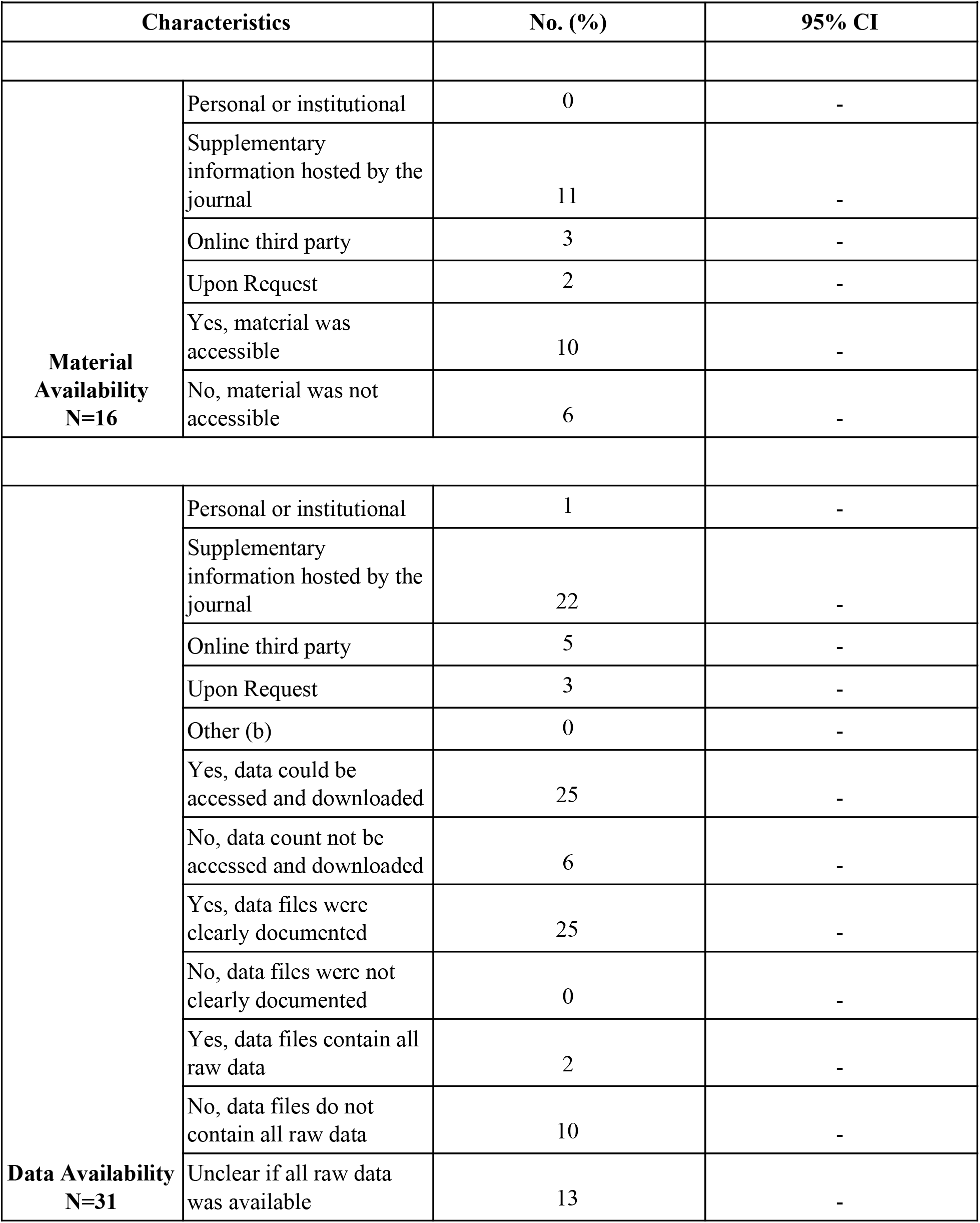
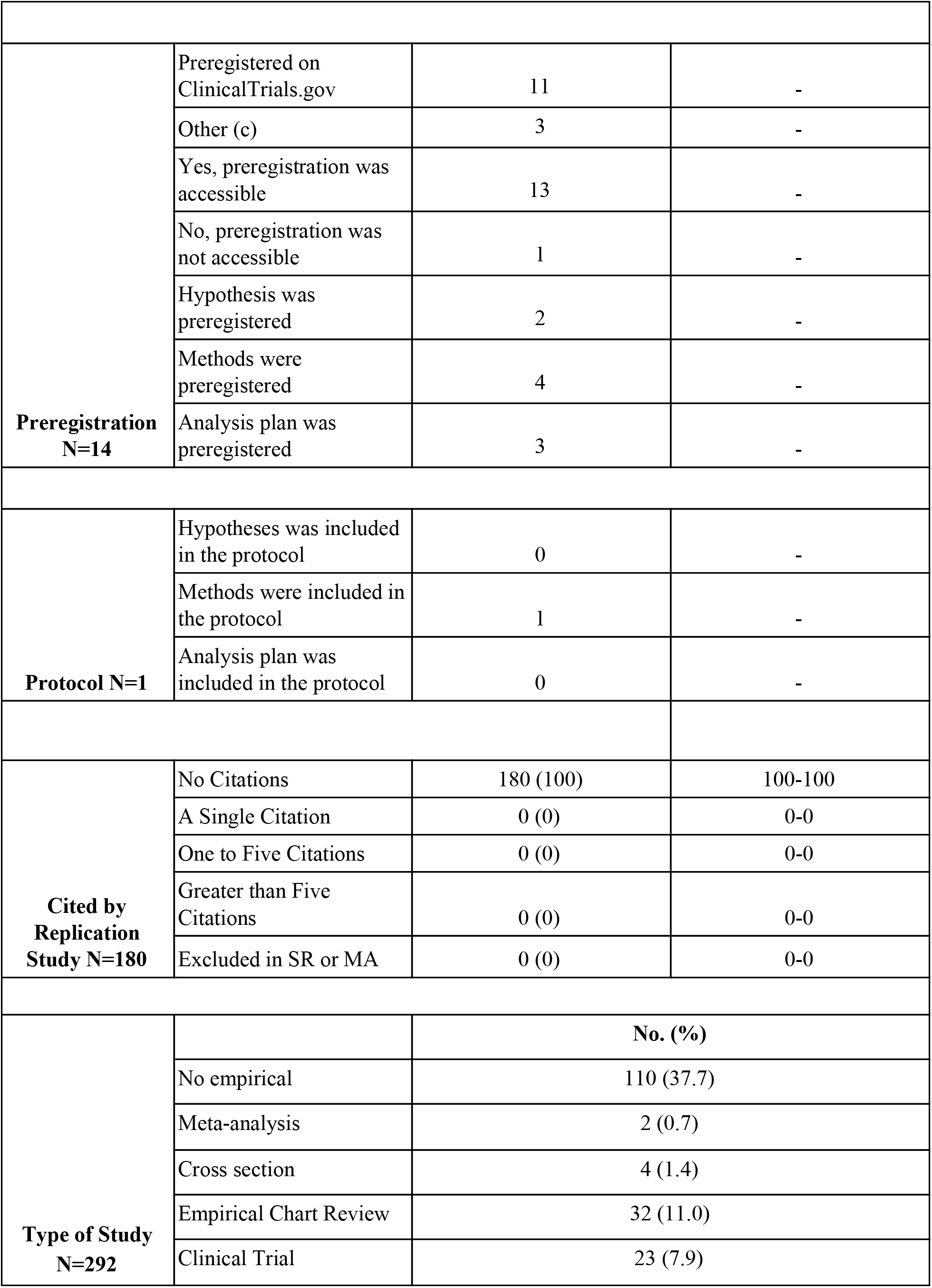
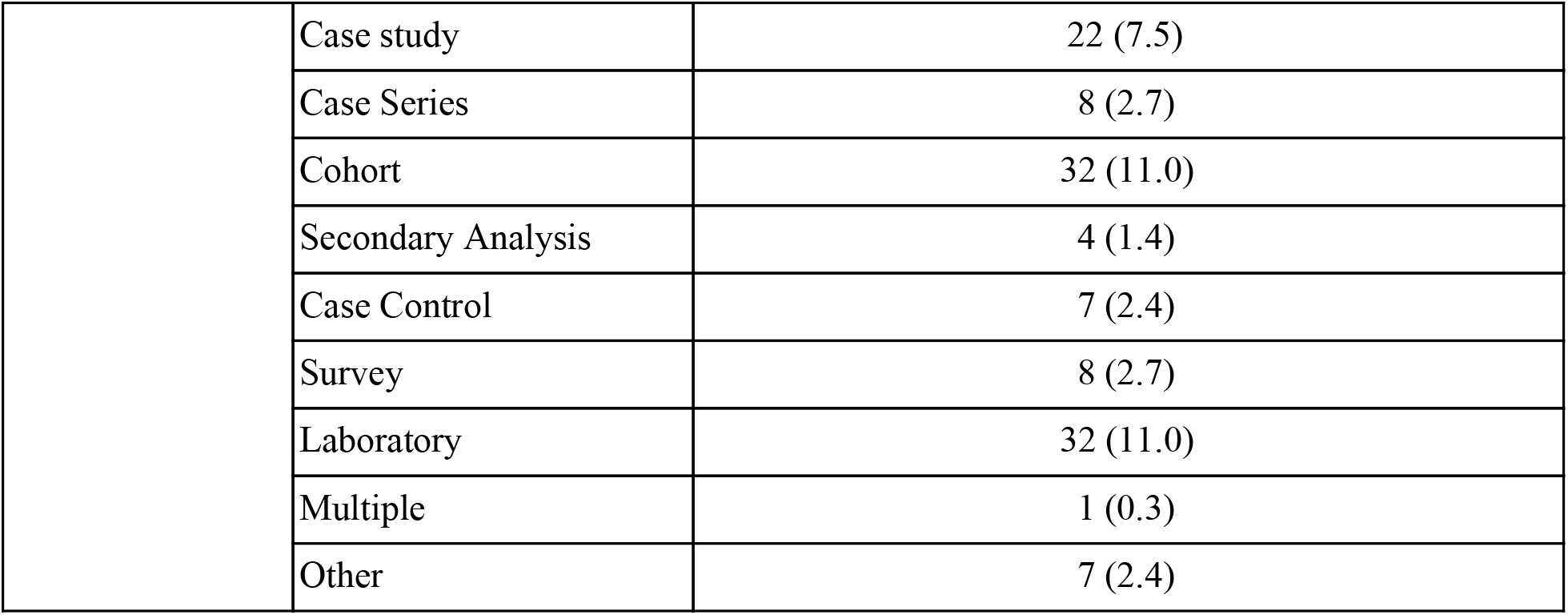
Additional Reproducibility Characteristics

**Figure 2: Proportion of Studies with Reproducibility Indicators**
| **Supplemental Table 1: Additional Reproducibility Characteristics** |  |  |  |
| --- | --- | --- | --- |
| **Characteristics** |  | **No. (%)** | **95% CI** |
| **Material Availability**<br>**N=16** | Personal or institutional | 0 | - |
|  | Supplementary information hosted by the journal | 11 | - |
|  | Online third party | 3 | - |
|  | Upon Request | 2 | - |
|  | Yes, material was accessible | 10 | - |
|  | No, material was not accessible | 6 | - |
| **Data Availability**<br>**N=31** | Personal or institutional | 1 | - |
|  | Supplementary information hosted by the journal | 22 | - |
|  | Online third party | 5 | - |
|  | Upon Request | 3 | - |
|  | Other (b) | 0 | - |
|  | Yes, data could be accessed and downloaded | 25 | - |
|  | No, data could not be accessed and downloaded | 6 | - |
|  | Yes, data files were clearly documented | 25 | - |
|  | No, data files were not clearly documented | 0 | - |
|  | Yes, data files contain all raw data | 2 | - |
|  | No, data files do not contain all raw data | 10 | - |
|  | Unclear if all raw data was available | 13 | - |
| **Preregistration**<br>**N=14** | Preregistered on ClinicalTrials.gov | 11 | - |
|  | Other (c) | 3 | - |
|  | Yes, preregistration was accessible | 13 | - |
|  | No, preregistration was not accessible | 1 | - |
|  | Hypothesis was preregistered | 2 | - |
|  | Methods were preregistered | 4 | - |
|  | Analysis plan was preregistered | 3 | - |
| **Protocol N=1** | Hypotheses was included in the protocol | 0 | - |
|  | Methods were included in the protocol | 1 | - |
|  | Analysis plan was included in the protocol | 0 | - |
| **Cited by**<br>**Replication**<br>**Study N=180** | No Citations | 180 (100) | 100-100 |
|  | A Single Citation | 0 (0) | 0-0 |
|  | One to Five Citations | 0 (0) | 0-0 |
|  | Greater than Five Citations | 0 (0) | 0-0 |
|  | Excluded in SR or MA | 0 (0) | 0-0 |
| **Type of Study**<br>**N=292** |  | **No. (%)** |  |
|  | No empirical | 110 (37.7) |  |
|  | Meta-analysis | 2 (0.7) |  |
|  | Cross section | 4 (1.4) |  |
|  | Empirical Chart Review | 32 (11.0) |  |
|  | Clinical Trial | 23 (7.9) |  |

|  |  |  |
| --- | --- | --- |
|  | Case study | 22 (7.5) |
|  | Case Series | 8 (2.7) |
|  | Cohort | 32 (11.0) |
|  | Secondary Analysis | 4 (1.4) |
|  | Case Control | 7 (2.4) |
|  | Survey | 8 (2.7) |
|  | Laboratory | 32 (11.0) |
|  | Multiple | 1 (0.3) |
|  | Other | 7 (2.4) |

